# Epigenetic differences between wild and cultivated grapevines highlight the contribution of DNA methylation during crop domestication

**DOI:** 10.1101/2023.10.12.562052

**Authors:** Alberto Rodriguez-Izquierdo, David Carrasco, Lakshay Anand, Roberta Magnani, Pablo Catarecha, Rosa Arroyo-Garcia, Carlos M. Rodriguez Lopez

**Affiliations:** Centro de Biotecnología y Genómica de Plantas (CBGP-INIA), CSIC - Universidad Politécnica de Madrid, Campus Montegancedo, Madrid, Spain; Environmental Epigenetics and Genetics Group (EEGG), Department of Horticulture, College of Agriculture, Food and environment, University of Kentucky, Lexington, KY, USA

**Keywords:** domestication, epigenomics, grapevine, epiGBS, wild, cultivated, methylation, epigenetic memory

## Abstract

The domestication process in grapevine facilitated the fixation of desired traits. The vegetative propagation of grapevines through cuttings has allowed for easier preservation of these genotypes compared to sexual reproduction. Nonetheless, even with vegetative propagation, different phenotypes often emerge within the same vineyard due to potential genetic somatic mutations in the genome. These mutations, however, are not the sole factors influencing phenotype. Alongside somatic variations, epigenetic variation has been proposed as pivotal player in regulating phenotypic variability acquired during domestication. The emergence of these epialleles might have significantly influenced grapevine domestication over time. This study aims to investigate the impact of the domestication process on the methylation patterns in cultivated grapevines. Reduced-representation bisulphite sequencing was conducted on 18 cultivated and wild accessions. Results revealed that cultivated grapevines exhibited higher methylation levels than their wild counterparts. Differential Methylation Analysis between wild and cultivated grapevines identified a total of 9955 differentially methylated cytosines, of which 78% where hypermethylated in cultivated grapevines. Functional analysis shows that core methylated genes (those consistently methylated in wild and cultivated accessions) are associated to stress response and terpenoid/isoprenoid metabolic processes. While genes presenting differential methylation are associated with proteins targeting to the peroxisome, ethylene regulation, histone modifications, and defense response. Additionally, our findings reveal that environmentally induced DNA methylation patterns are, at least partially, guided by the region of origin of wild grapevine accessions. Collectively, our results shed light on the pivotal roles that epialleles might have played throughout the domestication history of grapevines.

## Introduction

Domestication syndrome is a phenomenon observed in cultivated crops. This results in a suite of traits that distinguish them from their wild progenitors, including changes in morphology, physiology, and phenology that make them more amenable to cultivation. While the domestication syndrome has been well-documented in a handful of economically important seed propagated crops, thanks to the abundance of archaeobotanical, ecological and genetic information available for these species^1,2^, less is known about the domestication trajectories of vegetatively propagated crops^2^. One of the main advantages of vegetative propagation is that it allows for the preservation of desirable traits from one generation to the next. This is because when a plant is propagated vegetatively, the offspring is genetically identical to the parent plant^2–4^. This means that desirable traits such as disease resistance, yield, and flavor can be maintained over many generations. This contrasts with sexual reproduction, where traits can be lost or diluted through the process of genetic recombination. The type of propagation used during domestication can result in diametrically opposed domestication syndromes. For example, while the use of vegetative propagation has been shown to negatively affect the capacity for sexual reproduction via the accumulation of mutations in genes associated to flower development, self-fertilization, and seed development, which lead to the production of self-fertilized fruits, flowering asynchrony, and of less viable seeds^2,5,6^, crops domesticated by sexual reproduction, tend to present larger seeds, synchronic flowering and pollinator dependent fertillization^2^.

*V. vinifera* is a perennial woody liana belonging to the Vitaceae family, The species is divided into two different forms principally based on the sexuality of the plant and whether it is a cultivated or a wild form. Wild grapevines (*V. vinifera* ssp sylvestris), are commonly dioecious plants ^7^, and are naturally distributed across Asia and Europe. Cultivated grapevines (*V. vinifera* ssp vinifera), produce hermaphrodites flowers and is broadly cultivated across the world, both for grape production to be consumed as a fruit, and for winemaking, grape juice or other derived products ^8^.

Although the viticulture started at the Paleolithic age as a food source in Europe from wild accessions ^9^, there are evidences that the use of grapes by humans to produce wine started near to the seventh millennium BC ^10^, conditioning the domestication process in grapevine by selecting grapevine varieties producing a particular fruit quality and larger berries ^7,8^. It is believed that such selection occurred using vegetative propagation by cuttings to enhance the preservation of phenotypes of interest ^7,8,11^, which in turn had a negative effect on the crop tolerance to biotic and abiotic stresses. For example, populations of wild grapevines in North Africa and costal regions of Northern Spain shown better adaptation to salt stress than cultivated grapevines ^12,13^. While wild accessions from Germany, Iran and Georgia show higher resistance to mildew infections ^14–17^. Moreover, spite the use of vegetative reproduction to maintain a desired genotype, the use of asexual reproduction in grapevine has resulted in novel phenotypes appearing within the same variety^5^ and same vineyard ^18^. Such sports are frequently found in vegetatively propagated crops and often make up a significant portion of the cultivated varieties. Although a genetic basis is often presumed to be the reason for the noticeable differences in traits observed in sports, epigenetic modifications have also been proposed to play an important role^19-21^. Moreover, recent research have shown that DNA methylation epialleles can be used as an epimutation clock to enable the phylogenetic reconstruction of the recent history of vegetatively propagated plants^22^, highlighting their heritability and potential contribution to plant diversification. Nonetheless, our grasp of how these epigenetic modifications might have been utilized or unintentionally modified during the domestication process is currently at an early stage.

In this study, we use reduced representation bisulfite sequencing to characterize and compare the methylome of wild and cultivated grapevine accessions grown under common garden conditions, to explore if the domestication process has affected to methylome modelling in grapevine. We hypothesize that the combination of phenotype selection and vegetative propagation during grapevine’s domestication has resulted in methylomes characteristics of cultivated grapevines which are significantly different from those found in wild accessions. Moreover, we speculate that the epialleles observed in cultivated accessions could be associated to phenotypic traits traditionally associated to domesticated crops.

## Results

### Differences in global levels of DNA methylation between wild and domesticated grapevine genotypes

EpiGBS2 libraries yielded a total of 44.5 million reads with an average number 2.5 million of reads per sample (ranging from 1,106,659 to 8,249,031 reads). Bisulfite conversion efficiency showed on average 90% unmethylated cytosines converted to uracil’s. The average percentage of mappable reads per sample after de-multiplexing was 49%, ranging from 37% to 60%. This resulted in an average genome coverage of 1,5% (ranging between 0.7% and 2.6% (**Supplementary Table 1**), with reads distributed evenly across the whole genome (See **Figure 1A** for read distribution across chromosome 17 and **Supplementary File 1** for read distribution across all chromosomes).

**Figure 1:**
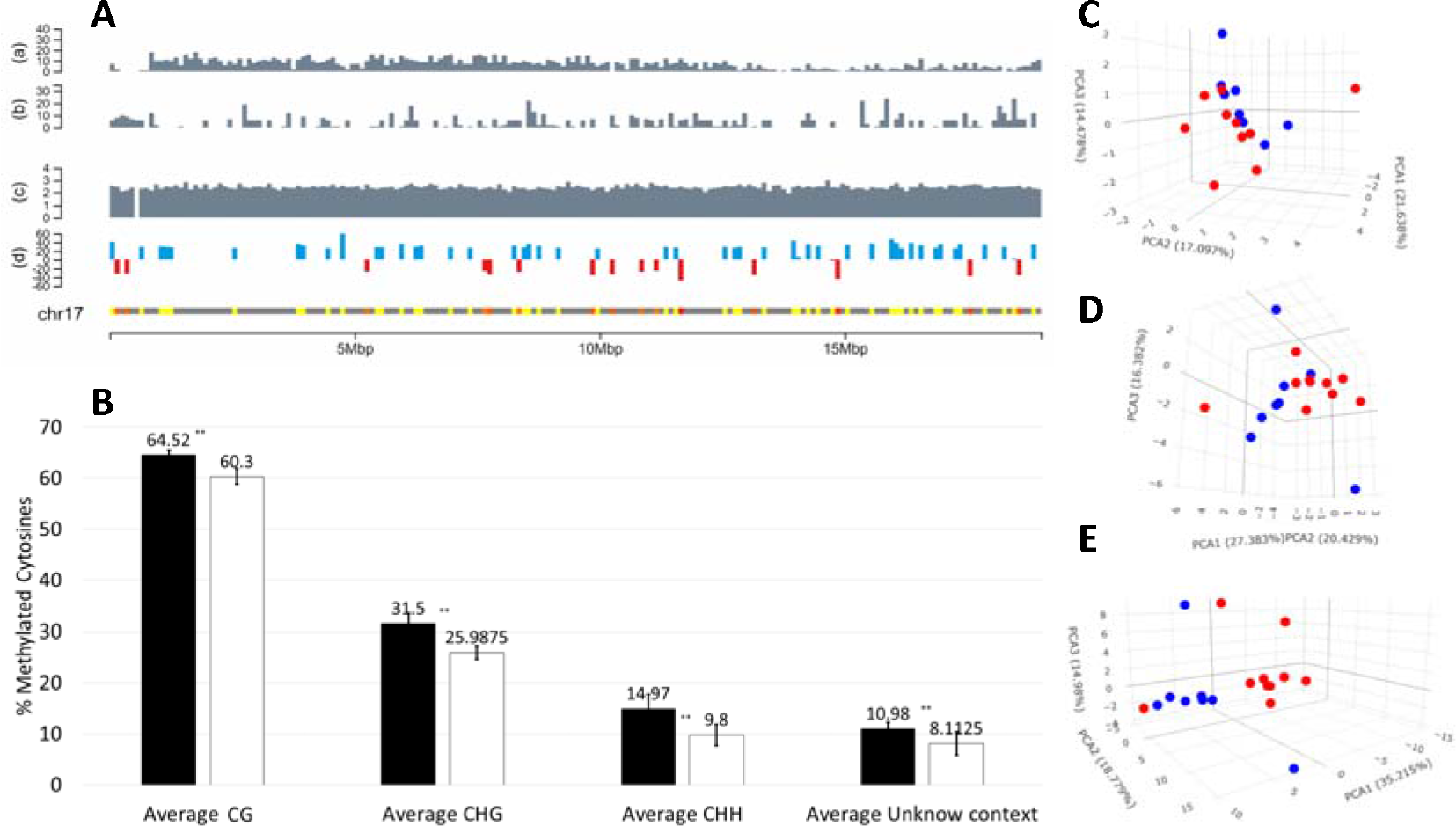
Analysis of differences in global levels of DNA methylation in cultivated and wild *V. vinifera* accessions. **A)** Visualization of genomic and epigenomic information for chromosome 17 of *Vitis vinifera using* 100,000 base pair windows. Vertical bars in panels (a) and (b) show the number of protein coding genes and transposable elements respectively per genomic window. Bars in panel (c) shows average sequencing depth per genomic window (Log 10 of calculated depth for sequenced bases). Panel (d) shows the average fold change in methylation in given window (blue and red bars indicate an average hypermethylated or hypomethylated window in cultivated vs wild accessions. For an interactive version of the figure containing all chromosomes see **Supplementary File 1**. Panels generated using ChromoMap R (Ref). **B)** Bars show the average percentage of methylation per sequence context (CG, CHG, CHH, and unknown) in cultivated (*V. vinifera ssp vinifera* (n = 10); black bars), and wild type (*V. vinifera* ssp *sylvestris* (n = 8); white bars) accessions. Error bars indicate the calculated Standard Deviation. ** T-student Test, p-value < 0.01. **C-E)** Multivariate analysis of percentage of methylation for all individual cytosine sequenced in cultivated and wild *V. vinifera* accessions. Principal Component analysis plots show results for methylation analysis results in the CG (C), CHG (D), and CHH (E) contexts. Blue and red circles represent cultivated and wild accessions respectively. PCs 1 to 3 represent 53, 64, 69% of the total measured variability in CG, CHG, and CHH contexts respectively.

Methylation calling showed that the CG context presented the highest level of cytosine methylation, followed by CHG and CHH context (**Figure 1B**). Cultivated varieties (CV) presented consistent significantly higher (T-student Test, p-value < 0.01) levels of DNA methylation than wild type accessions (WT) in all sequence contexts (**Figure 1B**). PCA plots built using the percentage of methylation for all sequenced cytosines as variables show that wild and cultivated form two different clusters separated mainly by PC1 in all sequence contexts (**Figure 1C-E**). Such observed separation between wild and cultivated accessions is particularly evident for the CHH context (**Figure 1E**).

### Identification of differentially methylated cytosines (DMCs) associated to domestication

Differential Methylation analysis identified a total of 9955 DMRs between wild and cultivated accessions evenly distributed across the genome (**Figure 1A** and **Supplementary File 1**). Of those, 7793 DMRs were hypermethylated and 2162 DMCs were hypomethylated in in wild cultivated grapevine accessions. The majority of both hyper and hypomethylated DMRs were found in the CHH context (77 and 69% respectively) (**Figure 2A**). From a gene feature context, DMRs were mainly found in intergenic regions **(Figure 2B)**. This is particularly evident in the CHH context, where 56 and 60% of hypermethylated and hypomethylated DMRs, respectively, were found in intergenic regions. The second most abundant genic feature presenting DMCs were introns, with percentages varying between 24 and 35% in hypermethylated DMRs, and 28 and 32% in hypomethylated DMRs, depending on the sequence context **(Figure 2B)**.

**Figure 2:**
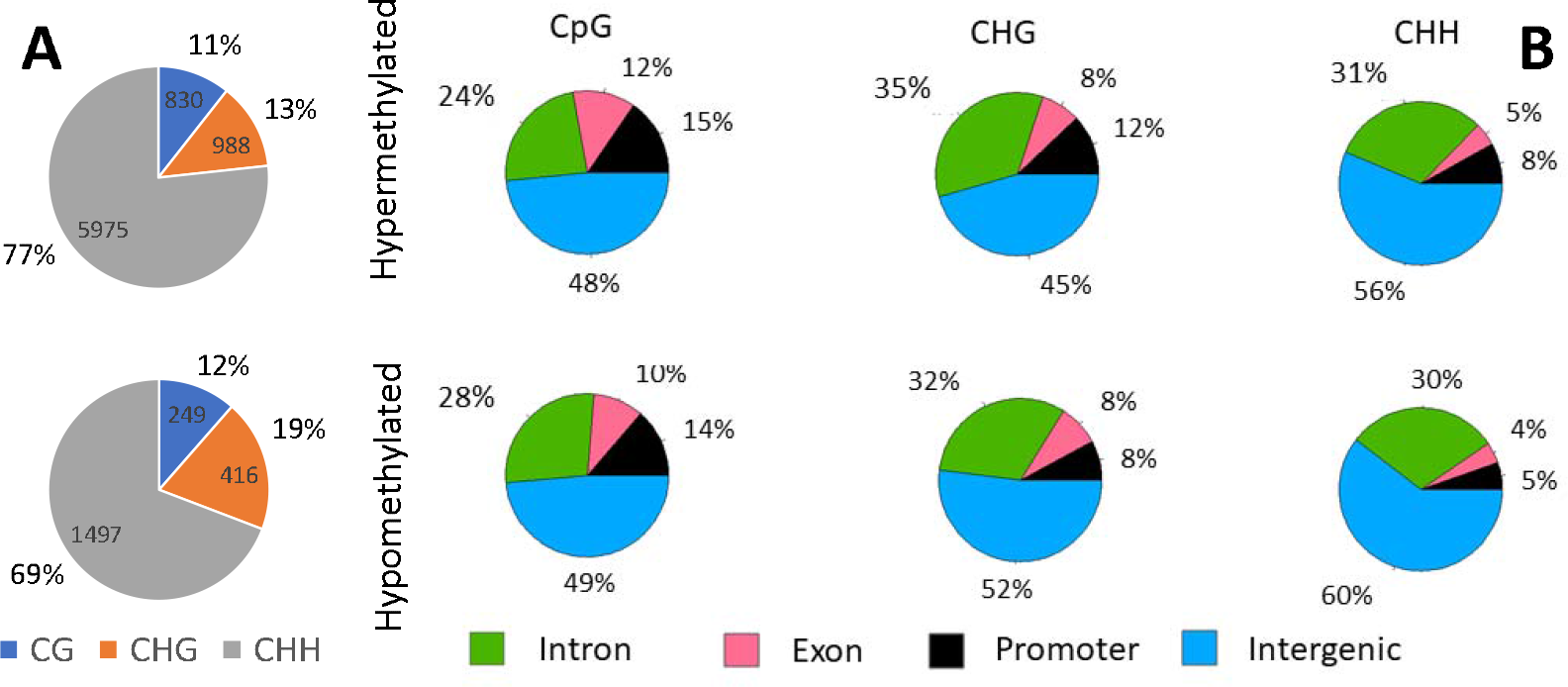
Identification of DMCs associated to grapevine’s domestication. Pie charts show **A)** the total number and percentage of hypermethylated (top pie chart) and hypomethylated DMCs identified in cultivated vines compared to wild accessions in each sequence context (CG, CHG and CHH); and **B)** the percentage of DMCs identified per genic feature and sequence context, in cultivated compared to wild type accessions.

### Analysis of epigenetic signals of provenance in wild type accessions

We then compared wild accessions to determine if an epigenetic signal associated to the location from where they were originally collected. To make sure that underlying genetic differences associated to geographic isolation between populations were not driving such epigenetic distance, we removed epialleles associated to SNPs using the epiDiverse-SNP pipeline. Epigenetic similarity analysis produced two separate clusters of wild accessions grouped by their provenance (**Figure 3**). One cluster contained all three accessions originally collected in the North of the Iberian Peninsula, in oceanic, continental and mountain climatic zones, while the second cluster contained all accession collected from the South of the Iberian Peninsula (Mediterranean climatic zone) (Figure 3) (see in **Supplementary Table 1** for metadata associated to each accession).

**Figure 3:**
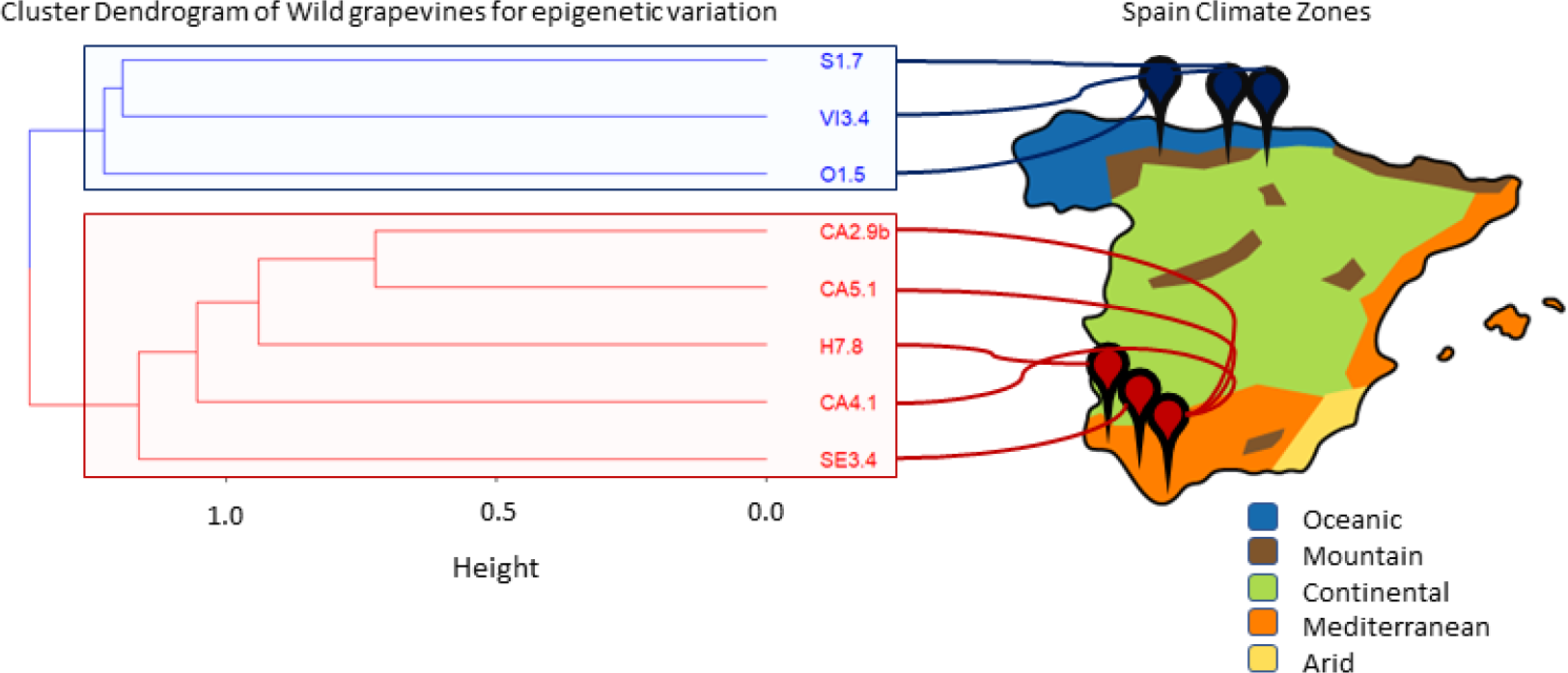
Effect of region of origin on the methylome of Iberian *Vitis vinifera sylvestris*. Analysis of the global differences in cytosine methylation between wild grapevine accessions grown under a common garden experiment but originally collected from different regions of the Iberian Peninsula. Analysis was performed using the epiDiverse-SNP pipeline to remove the effect of underlying genetic variation between wild grapevine populations. In red, the represented branches correspond to wild accessions belonging to the South of Spain, and in blue, they correspond to wild accessions coming from the North of Spain, placed approximately to the map of the Spanish Climate Zones.

### Analysis of domestication associated DMCs within genic features

Collectively epiGBS2 results generated reads overlapping with a total of 7174 genes. Of those, a total of 2854 (40%) genes were identified as genes that contained at least one methylated cytosine (Supplementary Table 2A). Methylated cytosines were mainly found in introns (66-80%), followed by exons (15-20%), and promoters (4-14%) **(Figure 4)** (Supplementary Table 2B). Genes containing methylated cytosines could be further divided into four groups, in order of abundance, 1. Genes presenting methylated cytosines both in wild and cultivated accessions (1883 genes) (core methylated genes (CMCs) hereafter); 2. Genes presenting CMCs and hypermethylated DMCs in cultivated compared to wild accessions (564 genes); 3. Genes presenting CMCs and hypomethylated DMCs in cultivated compared to wild accessions (252 genes); 4. genes presenting CMCs and both hypomethylated and hypermethylated DMCs (116 genes); 5. Genes presenting hypermethylated DMCs in cultivated compared to wild accessions (28 genes); and 6. Genes presenting hypomethylated DMCs in cultivated compared to wild accessions (11 genes). Functional analysis of the genes identified within each group revealed that CMCs are significantly associated with the regulation of cellular response to stress and isoprenoid/terpenoid processes. Cultivated grapevines hypermethylated genes were associated mainly to processes associated to protein targeting to peroxisomes and histone lysine demethylation, while hypermethylated genes in wild grapevines related to ethylene regulation processes and response to ozone. The remaining group (i.e., genes both presenting hyper and hypomethylated cytosines between in both types of accessions) presented GO terms related to defense response (Figure 4) (See **Supplementary Table 2C** for a complete list of GO terms in each group).

**Figure 4:**
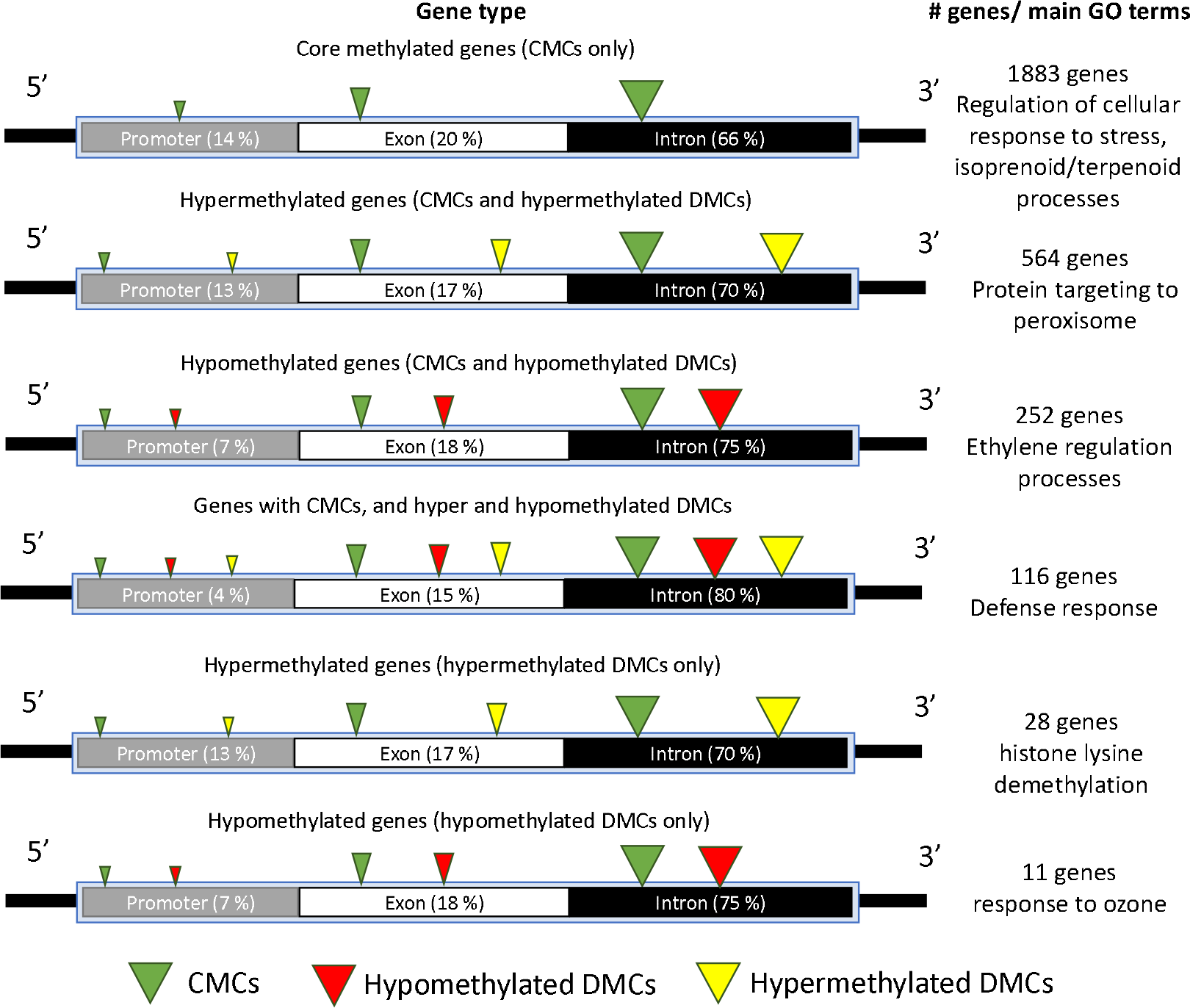
Schematic representation of methylated gene types in wild and cultivated grapevines. Boxes within gene models show the percentage of the total methylated cytosines in each gene group found in each genic context. Arrow heads color and size indicate the type of methylated cytosine found in each gene type (Core methylated cytosines (CMCs); Hypomethylated and hypermethylated cytosines in cultivated vs wild grapevine accessions) and the abundance of that type of methylation within that gene type, respectively. Right panel shows the number of identified genes for each group and their correspondent most representative GO terms.

## Discussion

While significant strides have been made in understanding the genetic underpinnings of crop domestication, there remains a comparative paucity of knowledge regarding the role of epigenetic mechanisms in this process. Epigenetics has emerged as a crucial regulator of various biological processes in both plants and animals. Recent studies have begun to hint at the potential involvement of epigenetic changes in the adaptation and phenotypic diversification of domesticated crops. However, a comprehensive understanding of how these epigenetic modifications may have been harnessed—or inadvertently altered—during the domestication process is still in its infancy. Studying the potential for epigenetic mechanisms contribution to domestication without genetic change^23-25^ could provide novel insights into the early stages of domestication and the selective pressures faced by ancestral agriculturists. At the same time, such studies will lay the foundation for the development of comprehensive models integrating plant adaptation to the environment through epigenetics mechanisms, facilitating their use for the development of novel cultivars more resilient to stress^26^.

### Epigenetic signal of geographic origin is stable and independent of genetic variation

Previous studies have shown that DNA methylation variability in plants can be attributed to three main factors: genetic (sequence) differences, environmental induction, and stochasticity (see Konate et al., 2020 for a recent example)^27^. Moreover, Xie et al., showed that cultivation method also contributes to environmentally induced epigenetic variability in grapevines^21^. Multiples studies have indicated the relationship between geographic origin and genetic differences in grapevine^7,28-30^, which would support the premise that genetic induced epigenetic differences should be observable between grapevines genotypes, independently of how long those accessions have been removed from their original source. However, it is not clear, if environmentally induced epigenetic variability is stable over time. To shed light over that question, we compared the methylome of wild grapevine accessions originally collected in different regions of the Iberian Peninsula, which have been maintained under the same growing conditions over 18 years. We observed that, after removing epialleles associated to SNPs, the methylomes of the studied accessions grouped according to the geographic precedence of the plants. Such clear differences among the wild accessions suggest that methylation patterns would be guided by their region of origin. Moreover, the maintenance of such epigenetic differences over such a protracted period of growth under the same environmental conditions, indicates that they might be fixed, and that they could, at least partially, explain phenotypic differences between phenotypes otherwise genetically very similar^22^.

### Grapevine domestic accessions present higher global levels of DNA methylation

Our DNA methylation analysis illuminated significant differences between cultivated and wild grapevines grown under the same conditions. An overall conspicuous increase in global methylation rates was observed in cultivated grapevines. This observation supports the hypothesis that cumulative methylation is a product of clonal propagation and domestication, as these processes foster selective epigenetic alterations that determine specific traits of interest^5,7^. In addition, Principal Component Analysis showed consistent differences between wild and cultivated grapevines, especially in the CHH and CHG contexts, a difference that was not as evident in the CG context. Such variations indicate that different factors might be influencing the methylation of cytosines in the CHH and CHG contexts, potentially outlining diverse selection pressures imposed by domestication for the different sequence context. Conversely, DMCs between wild and cultivated accessions were found evenly distributed across all chromosomes, suggesting that no specific epigenomic region has been under special selection. However, the observed prevalence of hypermethylated DMCs in cultivated grapevines across all contexts (CG, CHG, and CHH), underscores the postulation that cultivated grapevines accrue more epigenomic marks compared to their wild counterparts. This finding is in contradiction with recent studies showing that domestication induces a significant decrease in DNA methylation in rice. This is perhaps explained by the differences in the type of propagation used during the domestication of each species, which could have resulted in diametrically opposed domestication epigenetic syndromes. In fact, the historical use of vegetative propagation in cultivated grapevines^5,8^, has been shown to preserve environmentally induced epigenetic variability in vegetatively propagated perennials^31^. It is also tempting to speculate that the selection for hypermethylation during domestication, might have contributed to the appearance of genetic mutations leading to novel phenotypes^23^, since mutation ration is increased in methylated cytosines, as previously proposed for clone diversity in grapevines^32^.

### Domestication is associated with hypomethylation of intergenic regions and hypermethylation of genic features

As seen before^33^, in our study a large proportion of DMCs were found within intergenic regions, with hypermethylation being more prominent in wild than in cultivated accession. Nonetheless, 40% of the genes sequenced here presented methylated cytosines. Of these, 1883 (67%) presented only CMCs, i.e., cytosines which were consistently methylated both in wild and cultivated accessions, while the remaining 33% presented CMCs and or DMCs. CMCs and DMCs identified within genes were preferentially found within introns, followed by promoters and exons, irrespective of the sequence context. Contrary to what we observed in intergenic regions, DMC presenting genes were hypermethylated in cultivated grapevine accessions. This positional distribution of methylated cytosines around and within genes revealed different strategies in the methylation of genic features associated to the domestication process. Previous work has suggested that intergenic epialleles might be related to the regulation of long intergenic non-coding RNAs (lincRNAs), which are highly prevalent in the intergenic regions of plant genomes and are found to regulate essential biological processes^34^. It is also possible that the accumulation of methylation in intergenic regions could be related to silencing repeat elements or somatic mutations, which are a major driver of cultivated grapevine genome diversification^31^. In the context of plant promoters, methylation usually acts to repress gene transcription, thereby controlling the timing and spatial patterns of gene expression throughout development and in response to environmental stimuli^35^. In introns and exons, DNA methylation plays multifaceted roles. Within introns, it can influence alternative splicing, whereby different mRNA isoforms are generated from a single gene^36^. This can enhance the plant’s adaptive capability by enabling a diverse range of proteins to be produced. In exons, DNA methylation can impact gene-body methylation which is associated with increased gene expression in certain contexts, although the exact mechanism is not fully understood^36^. This intricate interplay between methylation and the genic landscape establishes a regulatory network that finely tunes gene expression and maintains genomic stability, underpinning the complexity and adaptability of plant life.

### Gene specific differential methylation associated to domestication is enriched in response to stress

Functional analysis of methylated genes showed that genes related to important agronomic traits exhibited significant DNA methylation level variation during grapevine domestication, particularly terms associated with stress response. This supports a large core of research linking epigenetic mechanisms in general, and DNA methylation in particular to plant response to stress^38^. Genes with differential methylation in the form of hypermethylation or hypomethylation between wild and cultivated grapevines were less abundant but still significant. The hypermethylated genes in cultivated grapevines were tied to protein targeting to peroxisomes and histone lysine demethylation. These processes are essential for cellular homeostasis and epigenetic regulation, suggesting that the domestication process may have enhanced or refined these functions in cultivated varieties. Interestingly, Histone H3-K4 demethylation, and DNA hypermethylation, have both been associated with gene expresión repression^39^. Conversely, genes hypermethylated in wild grapevines were found to relate to ethylene regulation processes and response to ozone. Ethylene is a critical hormone in plants, mediating various stress responses and developmental processes^40^. and ethylene responsive genes have been shown to change methylation status under abiotic stress^41^. The unique category of genes that showed both hyper and hypomethylated cytosines in both types of accessions, albeit being the smallest group, associated with defense response. Intriguingly, core methylated genes (CMCs) were also associated with stress response. This multimodal pattern of methylation during grapevine domestication suggests a complex regulation mechanism and might hint at genes that have retained some functionality from their wild origins, while also adapting new functionalities for the domesticated environment. Also, the conservation of methylation in the core methylated genes could suggest that the functions they support are essential and have remained unchanged between wild and cultivated grapevines.

In summary, our findings present compelling evidence that DNA methylation patterns have been significantly altered during the domestication of grapevines. The observed differential methylation patterns between wild and cultivated grapevine accessions offers fascinating insights into the potential roles that DNA methylation may play in the divergence between these two groups. While the varied associations of these methylation patterns to vital processes such as stress response, hormone regulation, and defense mechanisms underscore the importance of epigenetic regulation in shaping the evolutionary and developmental trajectories of domesticated species. Further studies, based on the analysis of complete methylomes, and focusing on the gene expression consequences of these methylation changes, would be valuable in elucidating the full spectrum of DNA methylation’s role in grapevine domestication.

## Materials and methods

### Experimental design

Single ortets from 10 *V. vinifera ssp vinifera* cultivars (Albillo Mayor, Allaren, Bocalilla, Brujidera, Espadeiro, Graciano, Heben, Jaen, Marfal and Zalema) and *vinifera* ssp *sylvestris* accessions originally collected from different places of Spain (see **Supplementary Table 1** for more information) (CA2.9b, CA4.1, CA5.1, H7.8, O1.5, S1.7, SE3.4 and VI3.4), and kept in a *in vivo* grapevine germoplasm bank located at IMIDRA (Instituto Madrileño de Investigación y Desarrollo Rural, Agrario y Alimentario, Alcalá de Henares, Madrid, Spain), were used to generated triplicate ramets from dormant wood cuttings. Cuttings were collected in winter, January 2021, at dormancy stage, from ortets planted on the same parcel. Cuttings were disinfected with tebuconazole and treated with rooting hormone (indole-butiric acid (IBA) 5 g/L), and then potted in individual containers (1.6 L truncated conic pots with drain sink) filled with potting mix 70 % peat / 20 % perlite / 10 % sand. All propagules were then placed under the same conditions (light 16h 21ºC - dark 8h 16ºC) in a single growth chamber, with all the cuttings distributed randomly along the growth chamber. After budbreak, the second and third fully open leaves were collected and immediately snap-frozen using liquid nitrogen and preserve at -80ºC until DNA extractions.

### DNA extraction and epiGBS protocol

Genomic DNA (gDNA) was extracted from all samples using the QIAGEN DNEasy Plant Mini Kit (Qiagen N. V., Hilden, Germany) following manufacturer’s instructions. DNA samples concentrations were determined using a Fragment Analyzer High Sensitivity DNA kit (Agilent). Sample concentration was standardized to 10ng/ul.

Reduced representation bisulphite sequencing (RRBS) libraries were prepared for all samples following the epiGBS2 protocol^42,43^ by digesting 100 ng of gDNA with restriction enzymes *Nsi*I and *Csp*6I (New England Biolabs, UK). Individually barcoded hemimethylated adapters, designed for the resulted restriction sites, were ligated to the resulting restriction products and amplified using PCR. Individual libraries generated from each sample were equimolarly mixed into two libraries which were sequenced using two Illumina HiSeq 2500 150bp paired-end runs by NovoGene USA.

### Bioinformatic analysis

Sequencing library quality was checked using FastQC v0.11.8^44^. A custom workflow was built to adapt the epiGBS workflow^42,43,45^ to our data. Firstly, demultiplexing was performed in order to ensure the structure of the adapters to identify the samples^43^, and a fastq-filter was performed using Stacks v2.55^46^. The demultiplexed sequences from the triplicates from each accession were pooled to form a unique sample. Pairedend sequences were merged using PEAR v0.9.6^47^. Alignment and methylation calling were performed with Bismark v0.23.0^48^ using the reference genome of *Vitis vinifera* L. PN40024 v4.1. Sequencing depth, coverage, and methylation differences between wild and cultivated accessions were visualized using ChromoMap R^49^.

Global differences in DNA methylation were visualized using hierarchical clustering and principal component analysis (PCA) performed on the calculated percentage of methylation in all methylated cytosines present in at least four of the accessions. Then the percentage of methylation for each cytosine was compared between cultivated and wild accessions in each context (CG, CHG and CHH (where H=A,T,C)) using T-student, considering significant differences when p-value < 0.01. Finally, differentially methylated cytosines (DMCs) were identified using the methylKit R package v1.16.1^50^. Cytosines were considered differentially methylated between wild and cultivated accessions when the observed difference in methylation was more than 25% and p-value < 0.01.

To determine if DNA methylation patterns associated to the geographic origin of wild accessions was present, we performed a comparative analysis following the premises of De Andrés et. al, (2012)^28^. For this, the methylation information gathered from wild accessions was filtered for epialleles associated to single nucleotide polymorphism. The remaining epialleles were used to cluster on wild grapevines in order to group it using epiDiverse - SNP pipeline (available at https://github.com/EpiDiverse/SNP).

Methylated cytosines within 1000 bp of protein coding genes identified using PN40024 v4.1 annotated genome to determine their position within genic features (promoters, introns, and exons). Genes presenting at least one methylated cytosine were deemed methylated. Then methylated genes were divided into 6 groups based on the type of methylation observed: 1. Core methylated genes, i.e., genes presenting unchanged methylated cytosines both in wild and cultivated accessions (CMCs); 2. Genes presenting CMCs and hypermethylated dfferentially methylated cytosines (DMCs) in cultivated compared to wild accessions; 3. Genes presenting CMCs and hypomethylated DMCs in cultivated compared to wild accessions; 4. genes presenting CMCs and both hypomethylated and hypermethylated DMCs; 5. Genes presenting hypermethylated DMCs in cultivated compared to wild accessions; and 6. Genes presenting hypomethylated DMCs in cultivated compared to wild accessions. Gene Ontology (GO) analysis was implemented with GOstats^51^ and rrvgo^52^ package in R, for each of these groups using all genes sequenced (i.e., presenting at least one read overlapping with a window of 1000 bp before and after the 5’ and 3’ UTRs respectively) as the gene universe. QuickGO Browser^53^ (GO version 2023-09-20) was used to generate the ancestor charts for the main GO terms in each group.

## Data availability

The datasets presented in this study can be found in the European Nucleotide Archive (ENA), accession number PRJEB55284

## Acknowledgements

This study was supported by the National Institute of Food and Agriculture, AFRI Competitive Grant Program Accession number 1018617, the National Institute of Food and Agriculture, United States Department of Agriculture, Hatch Program accession number 1020852, COST Action CA 17111 INTEGRAPE, supported by COST (European Cooperation in Science and Technology) and the Research and Science Ministry of Spain (project RTI2018-094470-R-C21).

## Conflict of interests

No conflict of interest declared.

## Contributions

ARI, RAG, and CMRL conceived the study. ARI, DC, RAG, and CMRL developed the experimental design for the study. RAG provided the plant accessions and metadata. ARI and RM performed the laboratory work. ARI, DC, and PC, performed the data analysis. ARI, DC, and LA performed data visualization. ARI and CMRL wrote the manuscript. All authors contributed to the article and approved the submitted version.

## Notes

### Competing Interest Statement

The authors have declared no competing interest.

